# A Suite of TMPRSS2 Assays for Screening Drug Repurposing Candidates as Potential Treatments of COVID-19

**DOI:** 10.1101/2022.02.04.479134

**Authors:** Jonathan H. Shrimp, John Janiszewski, Catherine Z. Chen, Miao Xu, Kelli M. Wilson, Stephen C. Kales, Philip E. Sanderson, Paul Shinn, Zina Itkin, Hui Guo, Min Shen, Carleen Klumpp-Thomas, Samuel G. Michael, Wei Zheng, Anton Simeonov, Matthew D. Hall

## Abstract

SARS-CoV-2 is the causative viral pathogen driving the COVID-19 pandemic that prompted an immediate global response to the development of vaccines and antiviral therapeutics. For antiviral therapeutics, drug repurposing allowed for rapid movement of existing clinical candidates and therapies into human clinical trials to be tested as COVID-19 therapies. One effective antiviral treatment strategy used early in symptom onset is to prevent viral entry. SARS-CoV-2 enters ACE2-expressing cells when the receptor-binding domain of the spike protein on the surface of SARS-CoV-2 binds to ACE2 followed by cleavage at two cut sites on the spike protein. TMPRSS2 has a protease domain capable of cleaving the two cut sites; therefore, a molecule capable of inhibiting the protease activity of TMPRSS2 could be a valuable antiviral therapy. Initially, we used a fluorogenic high-throughput screening assay for the biochemical screening of 6030 compounds in NCATS annotated libraries. Then, we developed an orthogonal biochemical assay that uses mass spectrometry detection of product formation to ensure that hits from the primary screen are not assay artifacts from the fluorescent detection of product formation. Finally, we assessed the hits from the biochemical screening in a cell-based SARS-CoV-2 pseudotyped particle entry assay. Of the six molecules advanced for further studies, two are approved drugs in Japan (camostat and nafamostat), two have entered clinical trials (PCI-27483 and otamixaban), while the other two molecules are peptidomimetic inhibitors of TMPRSS2 taken from the literature that have not advanced into clinical trials (compounds 92 and 114). This work demonstrates a suite of assays for the discovery and development of new inhibitors of TMPRSS2.

## Introduction

The severe acute respiratory syndrome-related coronavirus 2 (SARS-CoV-2) pandemic has driven the need for development of effective therapeutics to be used as prophylactics and as a treatment for infected patients. At the outset of the pandemic, there were no approved therapeutics for treating any coronaviruses, which led to the initiation of several human clinical trials on existing clinical candidates and approved drugs. Drug repurposing allows for drugs to be fast-tracked to the clinic for trials addressing a different disease than it was originally intended. An initial step in drug repurposing is the development of *in vitro* assays capable of generating efficacy data on the candidates for drug repurposing, which is critical for either supporting or denying their movement into human clinical trials. An example of this was remdesivir (GS-5734, Gilead Sciences Inc.) that had previously been in clinical trials for treating the Ebola virus but was repurposed as a SARS-CoV-2 treatment following its demonstration as a viral polymerase inhibitor within an *in vitro* experiment using live SARS-CoV-2 virus, leading to FDA approval.^1, 2^

Host cell targets have also received significant attention, partly because pharmacologic modulation should be invariable against viral variants/mutants.^3^ Host antiviral targets are often human proteins whose function has been co-opted and are essential to the viral replication cycle.^4^ Transmembrane protease serine 2 (TMPRSS2) is a human protein within the family of type-II transmembrane serine proteases (TTSP) expressed in epithelial cells of the human lungs, airways, and gastrointestinal tracts.^5, 6^ TMPRSS2 has become a therapeutic target due to its role in SARS-CoV-2 infection of a cell, and the existence of approved therapeutics able to inhibit its protease activity and be assessed for drug repurposing.^7^ A mechanism for SARS-CoV-2 entry includes binding of its surface glycoprotein spike (S) with the human cell-surface receptor angiotensin-converting enzyme 2 (ACE2), followed by cleavage at two sequences on the spike protein that can be done by human cell surface proteases, such as TMPRSS2, which causes a conformational change in spike resulting in membrane fusion and release of the viral RNA into the host cell.^8-10^ TMPRSS2 is considered the primary protease within the lung due to it having the highest co-expression with ACE2.^5^

We have reported an *in vitro* enzymatic assay using recombinant TMPRSS2 and a fluorogenic substrate, BOC-QAR-AMC, to demonstrate whether drug repurposing candidates, such as camostat and nafamostat, could inhibit the protease activity.^11^ Additionally, the enzymatic assay was used to evaluate hits identified from virtual screens.^12, 13^ Since our initial report on the development of an *in vitro* assay using recombinant TMPRSS2, several other *in vitro* assays have been developed. A cell-based *in vitro* assay using the fluorogenic substrate, BOC-QAR-AMC, in HEK293T cells overexpressing TMPRSS2 allows for quantification of proteolytic activity of TMPRSS2 and assessment of inhibitors within the cellular context compared to recombinant protein.^14^ Fluorescence resonance energy transfer (FRET)-based peptides mimicking the spike cleavage sites (S1/S2 and S2’) with recombinant TMPRSS2 demonstrated the capability of TMPRSS2 to cleave at both relevant spike cleavage sites.^15^ A split-luciferase *in vitro* assay using an engineered full-length human TMPRSS2 expressing a C-terminal HiBiT tag reported on both plasma membrane-associated and intracellular TMPRSS2 expression levels and demonstrated that homoharringtonine and halofuginone reduced total TMPRSS2 expression.^16^ Cell lines expressing TMPRSS2 (either endogenously or through transfection) have been used with pseudotyped particles or live virus assays to demonstrate the importance of TMPRSS2 in SARS-CoV-2 infection, and assess therapeutic candidates.^17, 18^ Altogether, assays on TMPRSS2 have been extensively used to assess therapeutic effectiveness.

Here, we report the development of a mass spectrometry-based detection assay using recombinant TMPRSS2 and an unlabeled peptide substrate that mimics the SARS-CoV-2 spike cleavage site, S2’. We demonstrate the suitability of the assay in 384-well plates, and its utility as an orthogonal assay capable of detecting false positive hits from the fluorogenic substrate assay. Additionally, the assay showed the ability of TMPRSS2 to cleave the spike cleavage site mimicked by the peptide, which agrees with results derived from use of FRET peptides within another study.^15^ Furthermore, we report the screening of inhibition of TMPRSS2 from several drug repurposing candidates within 6030 molecules in the NCATS drug libraries and an in-house compiled protease inhibitor library by using the fluorogenic peptide assay. Hits were further tested in the orthogonal mass spectrometry-based detection assay and a pseudotyped particle cell entry assay. The particles were pseudotyped with either the spike protein of the ‘wildtype’ (WA1 + D614G) or Delta strain (B.1.617.2), in Calu-3 human non-small cell lung cancer cells to demonstrate inhibition and cellular efficacy. Altogether, six molecules demonstrated activity in the biochemical and cell-based assays while remaining inactive in the counter assays. Two of these molecules, camostat and nafamostat, were the most potent inhibitors of TMPRSS2 and are both in human clinical trials as an antiviral against COVID-19.^19, 20^ Another two drug repurposing candidates, otamixaban and PCI-27483, demonstrated activity but have not entered human clinical trials for treatment of COVID-19. The final two molecules were identified from the literature as peptidomimetic inhibitors of TMPRSS2.^21^ Several other molecules were found to be weaker inhibitors. This work demonstrates an assay toolset for the discovery and development of new TMPRSS2 inhibitors.

## Results

### Assay design

Following our initial report on the development of a biochemical assay capable of monitoring TMPRSS2 activity and inhibition, we set out to conduct a drug repurposing screen.^11^ To briefly summarize the biochemical assay, yeast-expressed recombinant TMPRSS2 (aa 106-492, CUSAbio) was used along with a fluorogenic peptide substrate, BOC-QAR-AMC to monitor enzymatic activity via release of the caged 7-amino-4-methylcourmarin (AMC) fluorophore from the peptide at an emission wavelength of 440 nm (Figure 1). Buffer conditions were tested and optimized to include: 50 mM Tris pH 8, 150 mM NaCl and 0.01% Tween20. Using the peptide substrate at 10 μM, near its K_m_ of 33 μM, and TMPRSS2 at 175 nM, we achieved a signal-to-background (S:B) and Z’ of 3.2 and 0.86, respectively in a 1536-well plate. This demonstrated appropriate performance for this assay to be useful for quantitative high-throughput screening (qHTS).^22^

**Figure 1.**
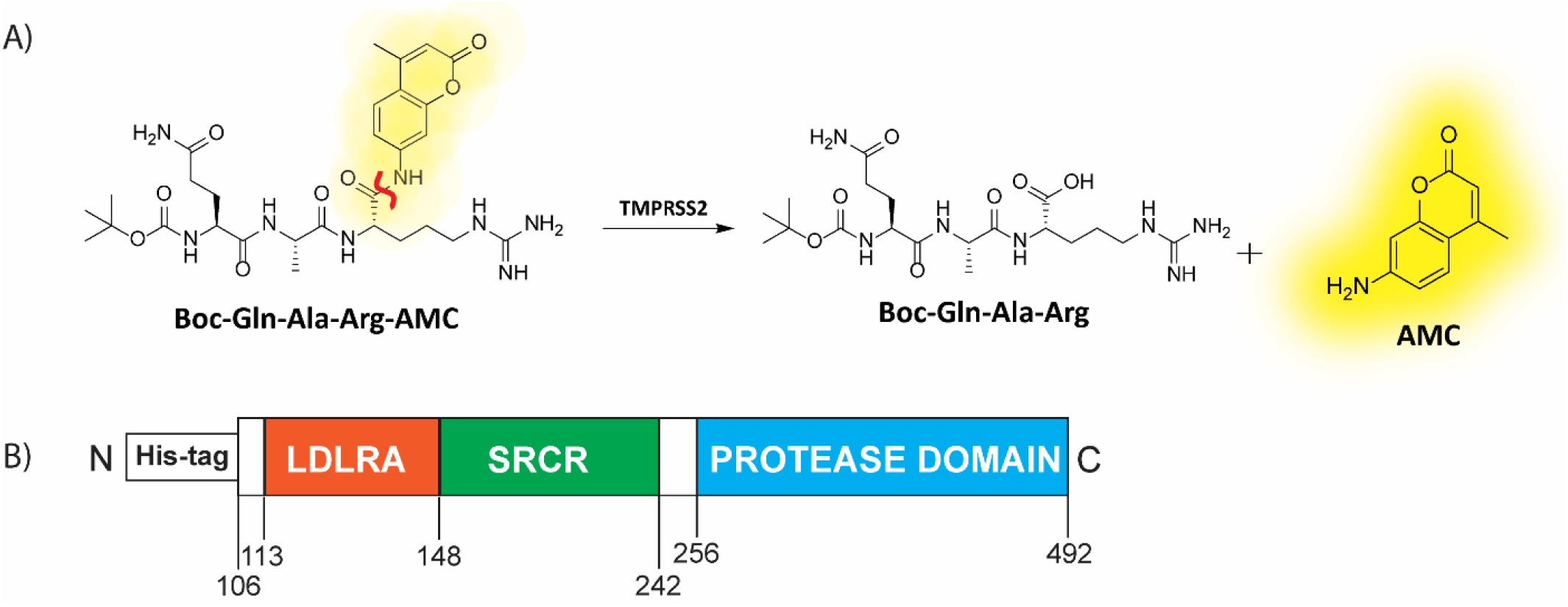
(A) Scheme displaying the enzymatic assay principle for the fluorogenic peptide substrate. The fluorogenic peptide substrate Boc-Gln-Ala-Arg-AMC has low fluorescence compared to the fluorescent 7-amino-4-methylcoumarin (AMC), which is released upon proteolytic cleavage. The scissile bond is indicated in red. (B) Schematic of the truncated yeast-expressed recombinant TMPRSS2 used in the fluorogenic assay, containing the low-density lipoprotein receptor A (LDLRA) domain, scavenger receptor cysteine-rich (SRCR) domain and protease domain.

### Drug repurposing qHTS of annotated small molecule libraries

Three compound libraries were selected for qHTS that are ideal for drug repurposing efforts: the NCATS Pharmaceutical Collection (NPC) contains all approved agents from USA, Europe, and Japan as of 2017 along with additional clinical candidates (2,678 compounds), the Mechanism Interrogation Plate (MIPE) is an oncology-focused library with a combination of approved and clinical agents (2,480 compounds), and an in-house compiled protease inhibitor library (PIL, 872 compounds) to screen against TMPRSS2, a total of 6,030 compounds (qHTS assay protocol shown in Table 1). Along with the potential for repurposing, the compounds in these libraries are annotated for published mechanisms of action. Each 1536-well plate contained 32 positive and 64 negative control wells to monitor assay performance. When screening the NPC library, the average plate Z’ was 0.76 with an average S:B of 2.39 (Figure 2A). The MIPE collection had an average plate Z’ of 0.72 with an average S:B of 2.33 (Figure 2B). The PIL collection had an average plate Z’ of 0.87 with an average S:B of 4.86 (Figure 2C).

**Table 1.** Detailed TMPRSS2 fluorogenic biochemical assay protocol for qHTS.

| Step # | Process | Notes |
| --- | --- | --- |
| 1 | 20 nL of peptide substrate dispensed into 1536-well plates. | Peptide (250x, dissolved in DMSO) was dispensed using an ECHO 655 acoustic dispenser (LabCyte) into a Corning 1536-well plate (Cat #3724). |
| 2 | Compounds and controls (20 nL) were pre-spotted into 1536-well plates. | Inhibitor (250x) or vehicle control (DMSO) was dispensed using an ECHO 655 acoustic dispenser (LabCyte). Compounds in dose-response were dispensed to columns 5-48 and controls dispensed into columns 1-4. |
| 3 | TMPRSS2 diluted in assay buffer dispensed into 1536-well plates. | TMPRSS2 (reconstituted in 50% glycerol at 8.75 μM, 50x) in assay buffer (50 mM Tris pH 8, 150 mM NaCl, 0.01% Tween20) was dispensed using a BioRAPTR (Beckman Coulter). Total reaction volume of 5 μL. |
| 4 | Incubated at RT for 1 hr | Final assay conditions were 10 μM peptide and 0.175 μM TMPRSS2 in assay buffer (50 mM Tris-HCl pH 8, 150 mM NaCl, 0.01% Tween20) |
| 5 | Read on PHERAstar FSX (BMG Labtech) | Fastest read settings, Fluorescence Intensity module: 340 nm excitation, 440 nm emission (Cat # 1601A2, BMG Labtech) |

**Figure 2.**
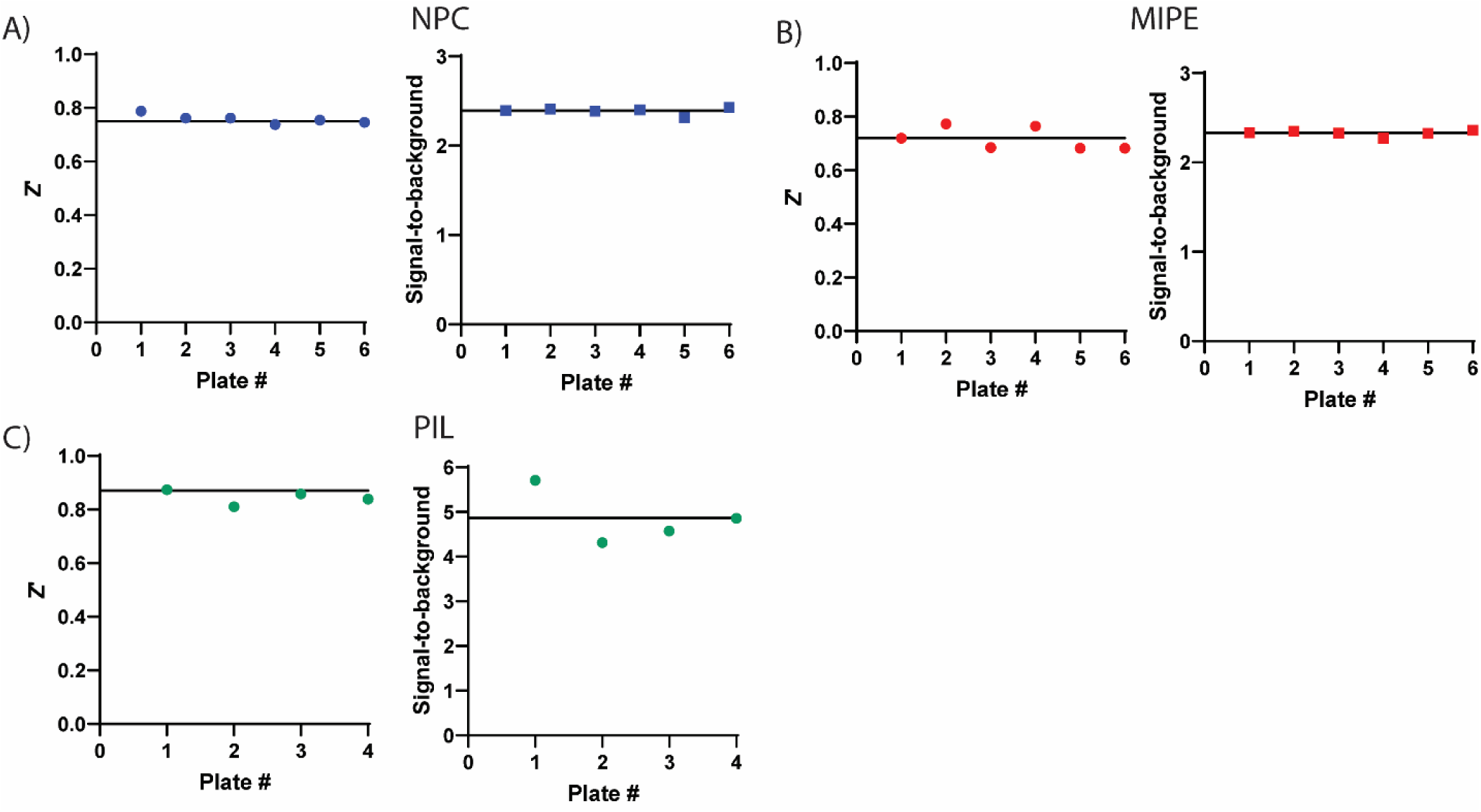
Assay performance for primary screening of the compound libraries in the TMPRSS2 fluorogenic biochemical activity assay. Z’ scores and signal-to-background values are plotted as data points for each plate. There were 32 positive and 64 negative control wells on each 1536-well plate. (A) NCATS Pharmaceutical Collection (NPC), 2,678 compounds, (B) Mechanism Interrogation Plate (MIPE) library, 2,480 compounds and (C) protease inhibitor library (PIL), 872 compounds. Black horizontal lines represent the mean values.

### Hit identification

The primary screen for NPC and MIPE were executed using compound concentrations of 40 μM, 8 μM and 1.6 μM, and hit selection criteria was set for those having a maximum response of ≥ -30% inhibition. A total of 14 compounds (0.5% hit rate) were identified as hits from the NPC library and 14 compounds (0.6% hit rate) were identified from the MIPE library. The primary screen for the PIL used compound concentrations of 40 μM, 10 μM, 2.5 μM, and 0.625 μM. Using the same hit selection criterion (max response ≥ -30% inhibition) yielded 37 active compounds (4.2% hit rate). Thus, from a screen of 6,030 compounds a total of 52 unique hits were identified, as there was redundancy from multiple hits presented between the libraries. Within the primary screen, 18,962 data points were generated. All qHTS data from the NPC and MIPE primary screens, and detailed protocol sheets, are available for download at the OpenData Portal website (https://opendata.ncats.nih.gov/covid19/). ^23^

### Hit confirmation

To confirm the hits identified from primary screening of the compound libraries, we tested each hit compound in triplicate dose-response in the primary fluorogenic biochemical assay using fresh stock. Compounds were plated in triplicate, 11-point, 1:3 dilution series using an ECHO 655 (LabCyte) acoustic dispenser. Of the 52 hits identified from the primary screening, 28 hits retained a max response inhibition of ≥ -30% when tested in triplicate dose-response. Next, the 28 hits were tested in a fluorescent counterassay, to detect false-positive hit compounds capable of quenching AMC fluorescence and mimicking inhibition, that involved the addition of compounds in dose-response to a buffer containing 1 μM AMC that approximates the estimated 10% enzymatic cleavage in the fluorogenic assay (i.e., starting concentration of substrate is 10 μM, counterassay protocol shown in Table 2). The counter-assay identified one compound (tannic acid) capable of quenching the AMC fluorescent signal ∼30%. Altogether, 27 compounds confirmed as inhibitors of TMPRSS2, which had ≥ -30% inhibition and ≤ -10% fluorescence quenching (Supplementary Table 1).

**Table 2.** Detailed fluorescence counterassay protocol for qHTS hits.

| Step # | Process | Notes |
| --- | --- | --- |
| 1 | 20 nL of 7-amino-4-methylcoumarin dispensed into 1536-well plates. | AMC (250x, dissolved in DMSO) was dispensed using an ECHO 655 acoustic dispenser (LabCyte) into a Corning 1536-well plate (Cat #3724). |
| 2 | Compounds and controls (20 nL) were pre-spotted into 1536-well plates. | Inhibitor (250x) or vehicle control (DMSO) was dispensed using an ECHO 655 acoustic dispenser (LabCyte). Compounds in dose-response were dispensed to columns 5-48 and controls dispensed into columns 1-4. |
| 3 | Assay buffer dispensed into 1536-well plates. | Assay buffer (50 mM Tris pH 8, 150 mM NaCl, 0.01% Tween20) was dispensed using a BioRAPTR (Beckman Coulter). Total reaction volume of 5 μL. |
| 4 | Incubated at RT for 5 min | Final assay conditions were 1 μM AMC with inhibitor or control (DMSO) in assay buffer (50 mM Tris-HCl pH 8, 150 mM NaCl, 0.01% Tween20). Columns 3 and 4 had 0.1 μM AMC with control (DMOS) in assay buffer. |
| 5 | Read on PHERAstar FSX (BMG Labtech) | Fastest read settings, Fluorescence Intensity module: 340 nm excitation, 440 nm emission (Cat # 1601A2, BMG Labtech) |

### Orthogonal mass spectrometry assay development

We next developed a biochemical assay using mass spectrometry and an unlabeled peptide substrate, with a peptide sequence from the S2’ site on the spike protein of SARS-CoV-2 (Figure 3A). Directly measuring an unlabeled peptide product, rather than a fluorescent product, removes the possibility of false positives from compound fluorescence interference.^24, 25^ The unlabeled peptide substrate sequence selected was Cbz-SKPSK**RS**FIED (scissile amide bond bolded), that yields cleavage products of Cbz-SKPSKR and SFIED by TMPRSS2 (Figure 3B). As the unlabeled peptide substrate sequence is derived from the SARS-CoV-2 spike protein TMPRSS2 cleavage site on the virus, it may be considered a more physiologically relevant approach to identifying and assessing TMPRSS2 inhibitors.

**Figure 3.**
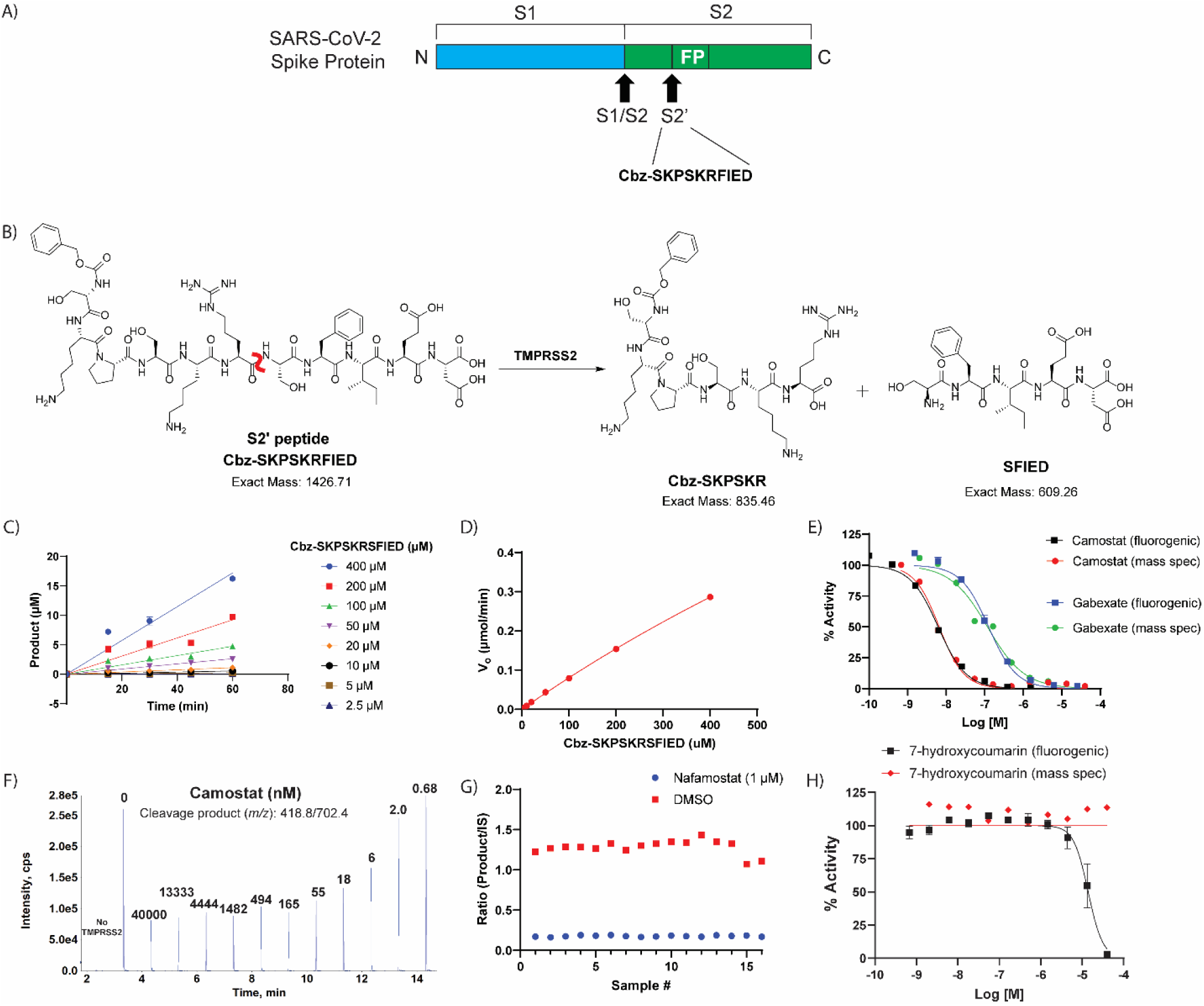
Label-free mass spectrometry biochemical assay. (A) Peptide derived from the known S2’ cleavage site of SARS-CoV-2 Spike (S) protein. (B) Scheme displaying the enzymatic assay principle for the unlabeled peptide substrate. The unlabeled peptide substrate, Cbz-SKPSKRFIED, is cleaved by TMPRSS2 to create two cleavage products, Cbz-SKPSKR and SFIED. The scissile bond is indicated in red. (C) Initial velocity (V_0_) was calculated for each substrate concentration by plotting product formation vs time. (D) The V_0_ for each concentration were plotted against the various substrate concentrations to obtain the V_max_ and K_m_. V_max_: 1.95 μmol/min and K_m_: estimated 2320 μM. (E) Comparison of TMPRSS2 fluorogenic detection and mass spectrometry detection assays by assessing dose-response inhibition by camostat and gabexate. (F) Mass spectrometry traces of the cleavage product (m/z: 418.8/702.4) showing that addition of camostat prevents product formation in dose-response. Each peak is labeled with the concentration of camostat (nM) for that condition. (G) Assay performance from the mass spectrometry detection assay when screening the hits identified from the primary screening. Z’ of 0.71 and S:B of 7.37. (H) Dose-response inhibition of 7-hydroxycoumarin against TMPRSS2 in both fluorogenic and mass spectrometry detection assays showing the fluorescent molecule as a false-positive hit in the fluorogenic assay, but not interfering with the mass spectrometry detection assay.

Linearity of detection for the spiked internal standard peptide (Cbz-SK-{^13^C_5_ ^15^N Pro}-SKR), which is the isotopically labeled version of the product peptide (Cbz-SKPSKR), was demonstrated through generation of a standard curve using concentrations from 1 nM to 2000 nM. To determine the K_m_ of Cbz-SKPSK**RS**FIED to guide assay substrate concentration,^26^ the TMPRSS2 concentration was held constant at 400 nM while substrate concentration varied from 2.5 μM to 400 μM. The enzymatic reaction was quenched by adding a quench solution (90:10 acetonitrile:water + 0.1% formic acid + 100 nM of the internal standard (Cbz-SK-{^13^C_5_^15^N Pro}-SKR), 1:1). The fragment peptides detected were the desired product (Cbz-SKPSKR, m/z 418.8/702.4) and the internal standard peptide (m/z 421.64/708.24), assay protocol shown in Table 3. Quantification of product formation was determined by observing the integrated area of the mass peak ratio between the desired product and internal standard. From the substrate titration experiment, the initial velocity was determined at each substrate concentration by plotting the amount of product formed against time (Figure 3C). Initial velocities were plotted against substrate concentrations (Figure 3D), indicating that the K_m_ was indeterminate as it yielded a value of 2320 μM, which is higher than the maximum substrate concentration in the assay. Due to the indeterminate K_m_, the assay conditions were set using an identical concentration of peptide substrate as the fluorogenic assay (10 μM). Under these conditions, the IC_50_ obtained from the mass spectrometry detection assay for camostat and gabexate were 6.6 nM (95% CI: 5.4, 8.2) and 119 nM (95% CI: 76.8, 181.3), respectively, which is statistically equivalent to that obtained from the fluorogenic assay, 6.2 nM (95% CI: 5.5, 7.0) and 130 nM (95% CI: 107.2, 147.5) (Figure 3E,F). Positive and negative control conditions were assessed in a 384-well plate format to demonstrate appropriate performance, which showed a S:B and Z′ of 7.37 and 0.71, respectively (Figure 3G).

**Table 3.**
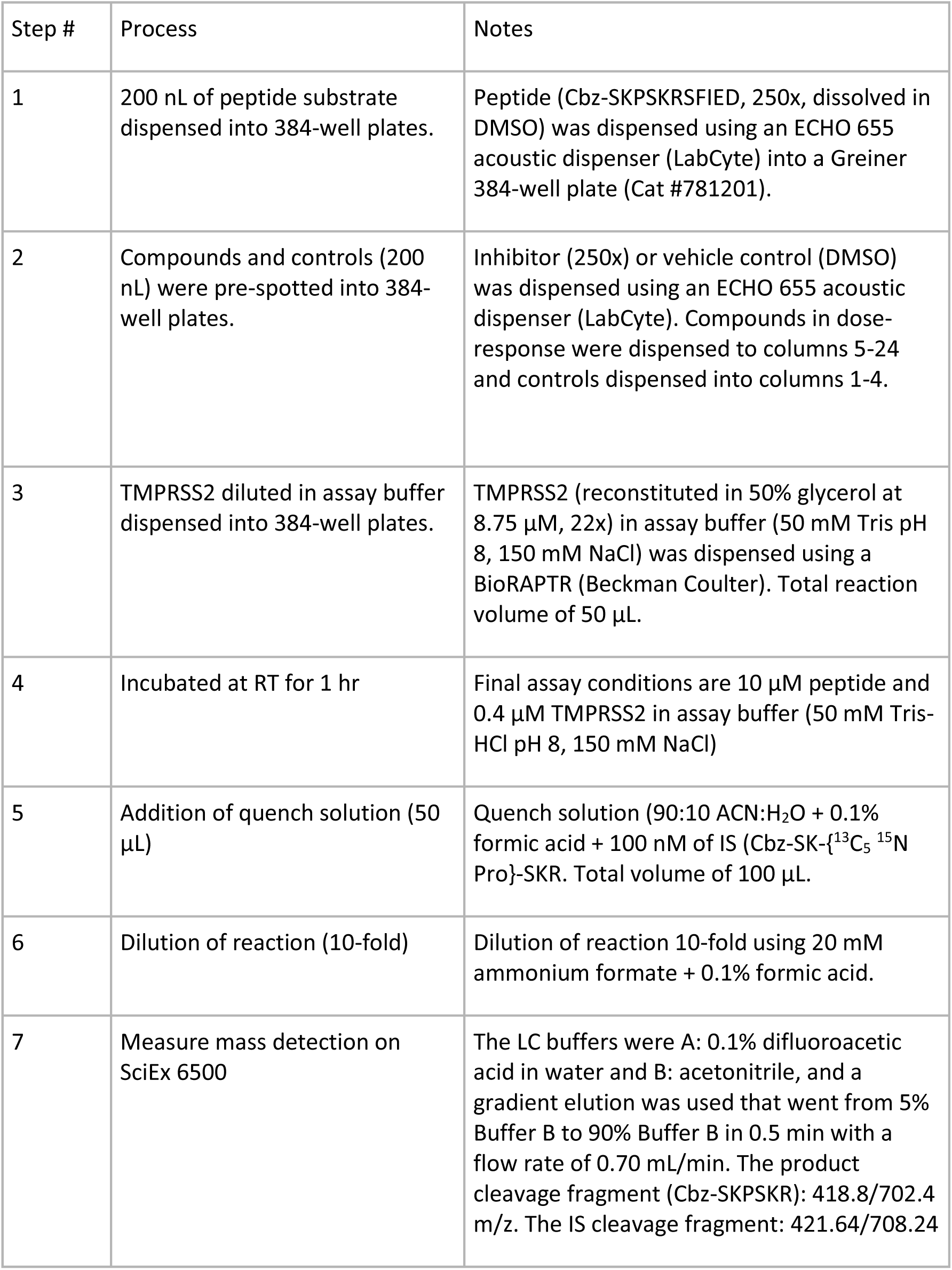
Detailed TMPRSS2 mass spectrometry detection biochemical assay.

| Step # | Process | Notes |
| --- | --- | --- |
| 1 | 200 nL of peptide substrate dispensed into 384-well plates. | Peptide (Cbz-SKPSKRSFIED, 250x, dissolved in DMSO) was dispensed using an ECHO 655 acoustic dispenser (LabCyte) into a Greiner 384-well plate (Cat #781201). |
| 2 | Compounds and controls (200 nL) were pre-spotted into 384-well plates. | Inhibitor (250x) or vehicle control (DMSO) was dispensed using an ECHO 655 acoustic dispenser (LabCyte). Compounds in dose-response were dispensed to columns 5-24 and controls dispensed into columns 1-4. |
| 3 | TMPRSS2 diluted in assay buffer dispensed into 384-well plates. | TMPRSS2 (reconstituted in 50% glycerol at 8.75 μM, 22x) in assay buffer (50 mM Tris pH 8, 150 mM NaCl) was dispensed using a BioRAPTR (Beckman Coulter). Total reaction volume of 50 μL. |
| 4 | Incubated at RT for 1 hr | Final assay conditions are 10 μM peptide and 0.4 μM TMPRSS2 in assay buffer (50 mM Tris-HCl pH 8, 150 mM NaCl) |
| 5 | Addition of quench solution (50 μL) | Quench solution (90:10 ACN:H<sub>2</sub>O + 0.1% formic acid + 100 nM of IS (Cbz-SK-{<sup>13</sup>C<sub>5</sub> <sup>15</sup>N Pro)-SKR. Total volume of 100 μL. |
| 6 | Dilution of reaction (10-fold) | Dilution of reaction 10-fold using 20 mM ammonium formate + 0.1% formic acid. |
| 7 | Measure mass detection on SciEx 6500 | The LC buffers were A: 0.1% difluoroacetic acid in water and B: acetonitrile, and a gradient elution was used that went from 5% Buffer B to 90% Buffer B in 0.5 min with a flow rate of 0.70 mL/min. The product cleavage fragment (Cbz-SKPSKR): 418.8/702.4 m/z. The IS cleavage fragment: 421.64/708.24 |

The 27 confirmed hits from the fluorogenic assay were tested in the mass spectrometry detection assay. One qHTS hit that failed to inhibit in the mass spectrometry detection assay was 7-hydroxycoumarin, a fluorescent molecule similar to 7-amino-4-methylcoumarin that is released upon cleavage of the fluorogenic substrate. Based on the fluorogenic assay, 7-hydroxycoumarin showed nearly full inhibition of TMPRSS2 at 40 μM; yet there was no inhibition detected within the mass spectrometry detection assay (Figure 3H). The cause for this discrepancy likely comes from 7-hydroxycoumarin being fluorescent; thereby, the endpoint fluorescent detection is overwhelmed by the amount of inhibitor fluorescence compared to the added cleaved product fluorescence. Since there is no increased fluorescence from the cleaved product at T60 (when quench solution is added) compared to T0, it appears that TMPRSS2 was fully inhibited with no cleaved product formation. This is an example of an assay interference false positive due to compound auto-fluorescence.^24^ The benefits of the mass spectrometry detection assay are twofold: 1) lower number of interference compounds identified, as demonstrated by 7-hydroxycoumarin and 2) capability of using a more physiologically relevant substrate that is unlabeled. Overall, 21 of the 27 compounds assessed for inhibition of TMPRSS2 showed activity. Unlike the assay interference compound, 7-hydroxycoumarin, the other five compounds that did not inhibit TMPRSS2 within the mass spectrometry detection assay was likely due to their weak activity within the fluorogenic assay.

### SARS-CoV-2-S Pseudotyped Particle Entry Assays

Finally, we demonstrated the effectiveness of these small molecules at preventing TMPRSS2-dependent viral entry into cells using a SARS-CoV-2 spike pseudotyped particle (PP) entry assay performed with the human airway epithelium cell line Calu-3, as previously described (Figure 4A,B).^18^ In addition to being the airway epithelium cell model, Calu-3 cells rely on serine proteases, like TMPRSS2, for viral entry when compared to other cell line models, which makes this cell line ideal for testing inhibitors of TMPRSS2.^17^ Two different pseudotyped particles with spike from different variants were used to assess TMPRSS2 inhibitors: one containing the Delta variant (B.1.617.2) spike protein that bears the mutations T19R, G142D, del156-157, R158G, L452R, T478K, D614G, P681R, D950N, and one containing the WA1 + D614G spike sequence. Notably, the Delta variant carries a P681R mutation in the S1/S2 cut site of the spike protein that may affect the cleavage activity by TMPRSS2; however, there are no mutations within the S2’ cut site. It’s unclear how the various mutations may affect the cleavage activity by TMPRSS2. A cytotoxicity counter assay was performed to assess whether hits were cytotoxic towards Calu-3 cells, which would produce a false-positive indicator of activity in the PP assay.

**Figure 4.**
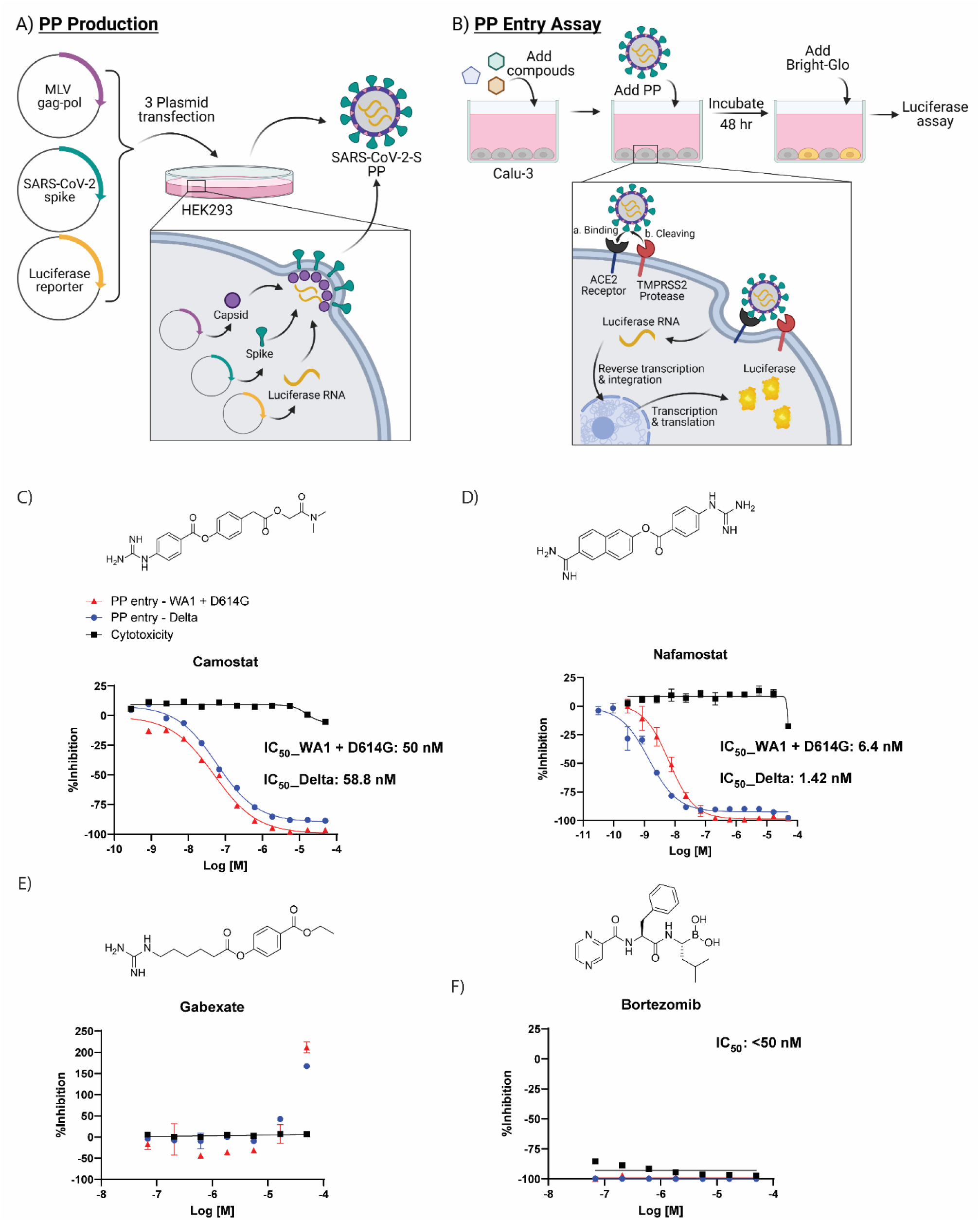
Cell-based pseudotyped particle assay. (A) Pseudotyped particle production. Three plasmids (pCMV-MLVgag-pol, pcDNA-SARS-CoV-2 spike, and pTG-Luciferase) are co-transfected into HEK-293T/17 cells. The plasmids express MLV core gag-pol polyprotein, SARS-CoV-2 spike glycoproteins, and luciferase RNAs, which together assemble into pseudotyped particles. (B) Pseudotyped particle assay scheme. Order of addition of reagents to Calu-3 cells is 1. compounds, 2. pseudotyped particles and 3. the Promega reagent Bright-Glo. Once cell entry of pseudotyped particle occurs, RNAs of pseudotyped particles are released into the cell, where they are reverse transcribed into DNAs, integrated into the genome and express luciferase reporter enzyme. The amount of luminescence following addition of Bright-Glo is proportional to the amount of pseudotyped particle entry into cells. Illustration was made with BioRender. Activity of clinically-approved inhibitors in prevention of pseudotyped particle entry using the Spike sequence from the Delta variant (blue), WA1 + D614G variant (red) and cytotoxicity counter-assay (black). The molecular structures and dose-response inhibition of particle entry by (C) camostat, (D) nafamostat, (E) gabexate, (F) bortezomib. The calculated concentrations required for 50% inhibition (IC_50_) are displayed in μM.

Compounds having confirmed inhibition of TMPRSS2 in both biochemical assays and in the cellular assay are listed in Table 4. Of the three most potent inhibitors of TMPRSS2 from the screening, camostat and nafamostat also demonstrated the greatest potency within the PP entry assay against the Delta pseudotyped particle with IC_50_ of 59 nM and 1.4 nM, respectively (Figure 4C,D). However, gabexate was inactive at inhibiting viral particle entry (Figure 4E). This result is in agreement with other studies reporting gabexate is inactive in cell-based assays testing TMPRSS2 inhibition.^27, 28^ Notably, bortezomib appeared to demonstrate a highly potent inhibition of viral entry (IC_50_ <50 nM); however, the cytotoxicity counter-assay revealed it was due to high cytotoxicity of the Calu-3 cells, which again demonstrates the importance and necessity of appropriate counterassays in designing an assay pipeline (Figure 4F). Additional evidence that 7-hydroxycoumarin is a false-positive identified from the fluorogenic assay is its lack of activity within the cell-based assay evaluating viral entry inhibition (Supplementary Table 1). The rest of the HTS hits had weak to no inhibition in the cell-based PP assay, most likely due to their already weak activity against TMPRSS2 from the biochemical assay. Notably, potency of the molecules was not significantly shifted between the two different types of pseudotyped particles from the two variants. This demonstrates the validity of drugging host targets such as TMPRSS2 to avoid viral escape of the antiviral therapy.

**Table 4.**
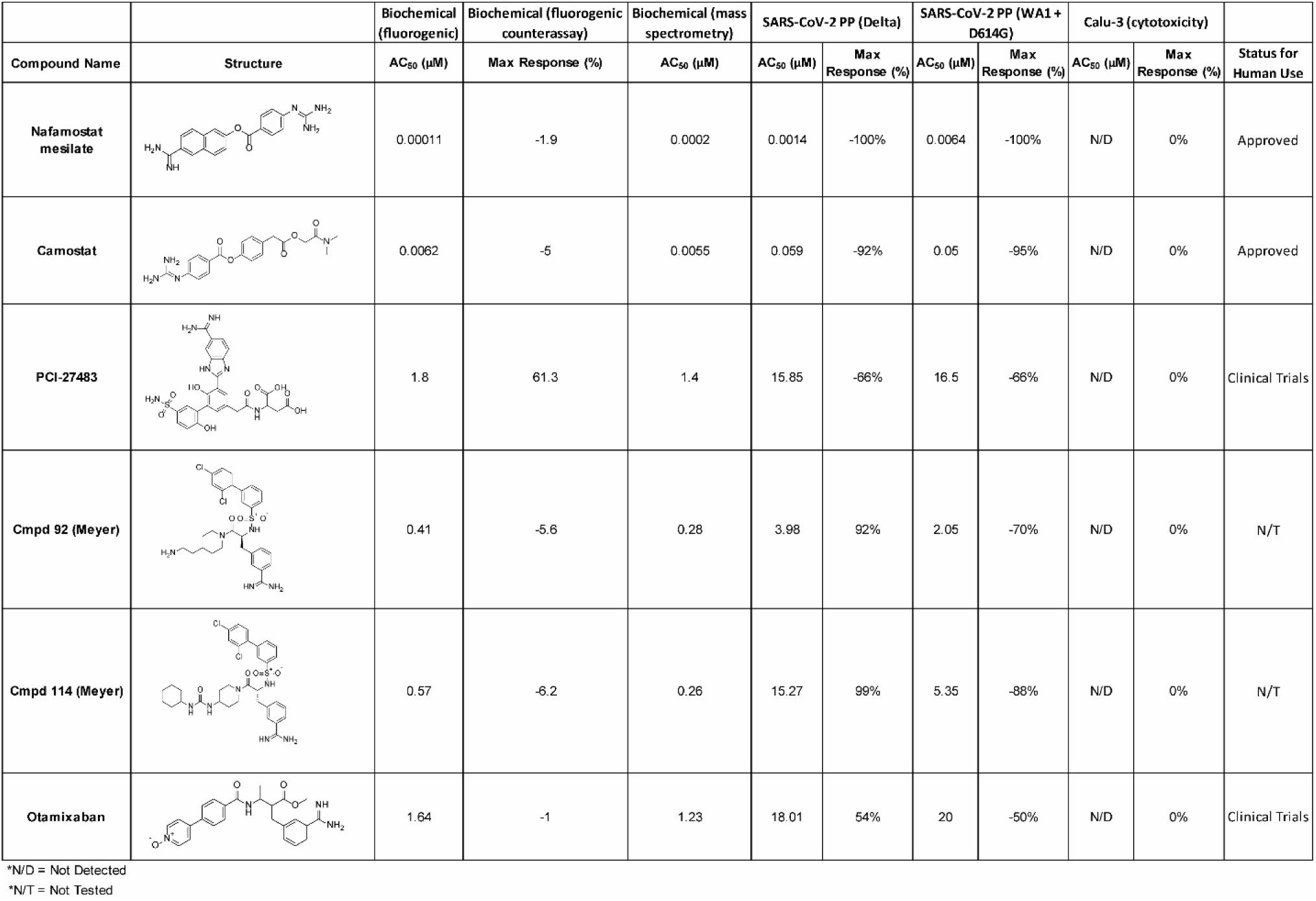
Compounds having confirmed inhibition of TMPRSS2 in both biochemical assays and in the cellular assay.

| Compound Name | Structure | Biochemical (fluorogenic) | Biochemical (fluorogenic counterassay) | Biochemical (mass spectrometry) | SARS-CoV-2 PP (Delta) |  | SARS-CoV-2 PP (WA1 + D614G) |  | Calu-3 (cytotoxicity) |  | Status for Human Use |
| --- | --- | --- | --- | --- | --- | --- | --- | --- | --- | --- | --- |
|  |  | AC<sub>50</sub> (μM) | Max Response (%) | AC<sub>50</sub> (μM) | AC<sub>50</sub> (μM) | Max Response (%) | AC<sub>50</sub> (μM) | Max Response (%) | AC<sub>50</sub> (μM) | Max Response (%) |  |
| Nafamostat mesilate |  | 0.00011 | -1.9 | 0.0002 | 0.0014 | -100% | 0.0064 | -100% | N/D | 0% | Approved |
| Camostat |  | 0.0062 | -5 | 0.0055 | 0.059 | -92% | 0.05 | -95% | N/D | 0% | Approved |
| PCI-27483 |  | 1.8 | 61.3 | 1.4 | 15.85 | -66% | 16.5 | -66% | N/D | 0% | Clinical Trials |
| Cmpd 92 (Meyer) |  | 0.41 | -5.6 | 0.28 | 3.98 | 92% | 2.05 | -70% | N/D | 0% | N/T |
| Cmpd 114 (Meyer) |  | 0.57 | -6.2 | 0.26 | 15.27 | 99% | 5.35 | -88% | N/D | 0% | N/T |
| Otamixaban |  | 1.64 | -1 | 1.23 | 18.01 | 54% | 20 | -50% | N/D | 0% | Clinical Trials |

\*N/D = Not Detected
\*N/T = Not Tested

## Discussion

Here we developed the first reported mass spectrometry-based detection assay for TMPRSS2 activity, and demonstrate a TMPRSS2 early drug discovery pipeline. To demonstrate proof-of-concept, we performed a HTS using our drug repurposing libraries.

The advantages to developing a mass spectrometry-based detection assay are twofold: 1) the unlabeled substrate can be more physiologically relevant leading to higher quality inhibitors being identified, 2) the unlabeled substrate avoids identification of a false positive that may arise in a fluorescence-based detection assay from inhibitor fluorescent quenching or compound fluorescence.^24^ For example, 7-hydroxycoumarin was identified as a hit within the fluorogenic assay. However, at the highest concentration tested of 40 μM, this molecule increases fluorescence to 1300x above the baseline level of fluorescence coming from the fluorogenic peptide substrate. Consequently, there is no increased fluorescence upon TMPRSS2 cleavage of the AMC from the fluorogenic peptide substrate; therefore, it instead appears that the enzyme was fully inhibited. The 7-hydroxycoumarin molecule did not show inhibition of TMPRSS2 in the mass spectrometry-based detection assay nor any inhibition of pseudotyped particle entry into Calu-3 cells.

Drug repurposing screening on *in vitro* assays yields initial proof-of-concept data that can be used to fast-track a drug to the clinic for trials addressing a different disease than it was originally intended. Our primary drug repurposing screen used recombinant TMPRSS2 with a fluorogenic peptide substrate with fluorescence detection. The screen included compounds from three in-house libraries totaling 6,030 compounds with many being drug repurposing candidates. Each compound was tested at multiple concentrations leading to generation of 18,962 data points. The primary screen yielded 52 hits. Upon assessing each of these compounds in an 11-point dose-response and in a fluorescence counter assay, only 27 compounds remained as confirmed hits. Each of these compounds were tested in dose-response for inhibition of TMPRSS2 in the mass spectrometry detection assay and in dose-response for inhibition of pseudotyped particle entry into Calu-3 cells.

Altogether, six compounds demonstrated inhibition of TMPRSS2 in biochemical and cellular assays. Those six compounds include two approved drugs (nafamostat and camostat, both approved in Japan to treat pancreatitis), two clinical candidates (PCI-27483 and otamixaban) and two literature compounds (compound 92 and compound 114).^21^

PCI-27483 was originally developed as a highly potent and selective inhibitor of the serine protease Factor VIIa (FVIIa) when in complex with tissue factor (TF) that has a normal physiological role in the initiation of the blood clotting cascade in response to an injured blood vessel wall.^29^ However, TF has been shown to be overexpressed in numerous primary tumors and is associated with tumor progression and worsened survival in cancer patients. This complex may contribute to tumor invasiveness by promoting cell migration and angiogenesis.^30^ Consequently, PCI-27483 entered clinical trials for patients with advanced pancreatic cancer in combination with gemcitabine. This trial has completed and though the combination was well tolerated, there was no demonstrated efficacy.^31^ PCI-27483 was first identified as a drug repurposing candidate as an inhibitor of TMPRSS2 by using a structure-based phylogenetic computational tool called 3DPhyloFold to identify structurally similar serine proteases, such as FVIIa, with known inhibitors that docked well to the TMPRSS2 structural model.^32^ Later, PCI-27483 was identified and confirmed to be an inhibitor of TMPRSS2 by NCATS scientists using a structural modeling and binding-site analysis of TMPRSS2 followed by a structure-based virtual screening and molecular docking approach.^12^ Though it has been demonstrated within *in vitro* studies as an inhibitor of TMPRSS2, it has yet to enter clinical trials as an antiviral treatment for COVID-19. The other clinical candidate, otamixaban, is an inhibitor of factor Xa (fXa), a critical serine protease in the blood coagulation cascade that catalyzes the conversion of prothrombin to thrombin.^33, 34^ Otamixaban entered clinical trials as an anticoagulant for patients with non-ST-segment elevation acute coronary syndromes (NSTE-ACS) against a control arm using unfractionated heparin plus eptifibatide. Otamixaban was shown to be safe but lack efficacy relative to the control arm.^35^ Otamixaban was first identified as a drug repurposing candidate as an inhibitor of TMPRSS2 within a virtual screen.^36^ Corroborating data came from the Noe lab and our group that included computational, biochemical and cellular data to demonstrate its inhibition of TMPRSS2.^12, 13^ It has yet to enter clinical trials as an antiviral treatment for COVID-19.

The two literature compounds, **92** and **114**, have sulfonylated 3-amindinophenylalanylamide core structures that were among the initial synthetic inhibitors of TMPRSS2.^21^ The Steinmetzer lab, developed a fluorescence based assay using a fluorogenic peptide substrate used with recombinant TMPRSS2 to identify **92** and **114** as potent inhibitors with a K_i_ of 0.9 nM and 5 nM, respectively. Both were shown to be not cytotoxic within Calu-3 cells. Additionally, **92** significantly reduced the replication of H1N1 and H3N2 influenza virus strains in Calu-3 cells. However, no further studies had been done with respect to SARS-CoV-2 with these compounds. We demonstrate inhibition of TMPRSS2 with an IC_50_ of ∼300 nM for both compounds and blocking of entry into Calu-3 cells using pseudotyped particles with SARS-CoV-2 spike protein with an IC_50_ of 4 μM and 15 μM for **92** and **114**, respectively.

## Methods

### Reagents

Recombinant Human TMPRSS2 protein expressed from yeast (human TMPRSS2 residues 106−492, N-terminal 6x His-tag) (cat.# CSB-YP023924HU) was acquired from Cusabio. The fluorogenic peptide substrate, Boc-QAR-AMC. HCl was obtained from Bachem (cat.# I-1550). The unlabeled peptide substrate, Cbz-SKPSKRSFIED was custom ordered from LifeTein.

### Fluorogenic Peptide Screening Protocol 1536-Well Plate

To a 1536-well black plate was added Boc-QAR-AMC substrate (2.5 mM, 20 nL, 250x) and inhibitor (20 nL, 250x) using an ECHO 655 acoustic dispenser (LabCyte). To that was dispensed TMPRSS2 (reconstituted in 50% glycerol at 8.75 μM, 50x) in assay buffer (50 mM Tris pH 8, 150 mM NaCl, 0.01% Tween20) using a BioRAPTR (Beckman Coulter) to give a total reaction volume of 5 μL. Following 1 h of incubation at RT, detection was done using the PHERAstar with 340 nm excitation and 440 nm emission (Table 1).

### Fluorescence Counter Assay 1536-Well Plate

To a 1536-well black plate was added 7-amino-4-methylcoumarin (0.25 mM, 20 nL, 250x) and inhibitor (20 nL, 250x) to columns 5-48 using an ECHO 655 acoustic dispenser (LabCyte). To the control columns 1-2 was added 7-amino-4-methylcoumarin (0.25 mM, 20 nL, 250x) and DMSO (20 nL), and to the control columns 3-4 was added 7-amino-4-methylcoumarin (0.025 mM, 20 nL, 250x) and DMSO (20 nL). Assay buffer (50 mM Tris pH 8, 150 mM NaCl, 0.01% Tween20) was added to the full plate to give a total reaction volume of 5 μL. Detection was done using the PHERAstar with 340 nm excitation and 440 nm emission. Fluorescence was normalized relative to a negative control containing 7-amino-4-methylcoumarin at 0.1 μM (−100% activity, low fluorescence) and a positive control containing 7-amino-4-methylcoumarin at 1 μM (0% activity, high fluorescence). An inhibitor causing fluorescence quenching would be identified as having a concentration-dependent decrease on AMC fluorescence resulting in a <0% activity (Table 2).

### Mass Spectrometry-based Assay 384-well plate

To a 384-well plate (Greiner 781201) was added Cbz-SKPSKRSFIED substrate (2.5 mM, 200 nL, 250x) and inhibitor (200 nL, 250x) using an ECHO 655 acoustic dispenser (LabCyte). To that was dispensed TMPRSS2 (reconstituted in 50% glycerol at 8.75 μM, 22x) in assay buffer (50 mM Tris pH 8, 150 mM NaCl) using a BioRAPTR (Beckman Coulter) to give a total reaction volume of 50 μL. Following 1 h of incubation at RT, the reaction was quenched by adding 90:10 ACN:H_2_O + 0.1% formic acid + 100 nM of internal standard (Cbz-SK-{^13^C_5_^15^N Pro}-SKR, 50 μL) to give a total volume of 100 μL. Samples were further diluted 10-fold using 20 mM ammonium formate + 0.1% formic acid prior to injection into a SciEx6500, and the data were analyzed using Analyst Software. An LS-1 autosampler (Sound Analytics, Niantic, CT) injected samples to the 2×20mm Halo C18 (Mac-Mod) column for LC separation. The LC buffers were A: 0.1% difluoroacetic acid in water and B: acetonitrile, and a gradient elution was used that went from 5% Buffer B to 90% Buffer B in 0.5 min with a flow rate of 0.70 mL/min. The cleavage fragment (Cbz-SKPSKR): 418.8/702.4 m/z was quantified to asses TMPRSS2 activity (Table 3).

### Data Process and Analysis

To determine compound activity in the assay, the concentration−response data for each sample was plotted and modeled by a four-parameter logistic fit yielding IC_50_ and maximal response values. Raw plate reads for each titration point were first normalized relative to a positive control containing nafamostat at 1 μM (0% activity, full inhibition) and a negative control containing DMSO-only wells (100% activity, basal activity). Data normalization, visualization, and curve fitting were performed using Prism (GraphPad, San Diego, CA).

### SARS-CoV-2-S Pseudotyped Particle Entry Assays

A SARS-CoV-2 Spike pseudotyped particle (PP) entry assay was performed as previously described (Chen et. al.) In short, 40,000 cells/well Calu3 cells (catalog HTB-55, ATCC) were seeded in white, solid bottom 96-well microplates (Greiner BioOne) in 100 μL/well media and incubated at 37 °C with 5% CO_2_ overnight (∼16 h). Then, the supernatant was removed, and compounds were added as 50 μL/well, 2X solutions in media. Cells were incubated with compounds for 1 h at 37 °C with 5% CO2, before 50 μL/well of SARS-CoV-2-S pseudotyped particles (PP) was added. We used the PP pseudotyped SARS-CoV-2 spike from the Delta variant B.1.617.2 (catalog 21H13, Codex Biosolutions) with a C-terminal 19 amino acid deletion using the murine leukemia virus system as previously described (Millet et. al.). The plates were then spinoculated by centrifugation at 1500 rpm (453g) for 45 min and incubated for 48 h at 37 °C with 5% CO_2_ to allow the cell entry of PP and the expression of the luciferase reporter. After the incubation, the supernatant was removed, and 20μL/well of the Bright-Glo Luciferase detection reagent (catalog E2620, Promega) was added to assay plates and incubated for 5 min at room temperature. The luminescence signal was measured using a PHERAStar plate reader (BMG Labtech). The data were normalized with wells containing SARS-CoV-2-S PPs as 0% and wells containing control bald PP as -100%. A cytotoxicity counter screen was performed using the same protocol without the addition of PP, with an adenosine triphosphate (ATP) content assay kit (catalog 6016949, Promega).

## Supporting Information

The Supporting Information is available free of charge at _.

Supplementary Table 1: A list of the 27 compounds as confirmed hits following screening in the fluorogenic assay and fluorescence counterassay.

## Acknowledgement

The authors acknowledge James Inglese for helpful discussions on enzyme kinetics. This work was supported by the National Center for Advancing Translational Sciences, Division of Preclinical Innovation.

